# Widespread translational control regulates retinal development in mouse

**DOI:** 10.1101/2021.03.02.433656

**Authors:** Kaining Chen, Congying Chen, Huihui Li, Jiaqi Yang, Mengqing Xiang, Hongwei Wang, Zhi Xie

## Abstract

Retinal development is tightly regulated to ensure the generation of appropriate cell types and the assembly of functional neuronal circuitry. Despite remarkable advances that have been made in understanding the regulation of gene expression during retinal development, how translational regulation guides retinogenesis is less understood. Here, we conduct a comprehensive translatome and transcriptome survey to the mouse retinogenesis from the embryonic to the adult stages. We discover thousands of genes that have dynamic changes at the translational level and pervasive translational regulation in a developmental stage-specific manner with specific biological functions. We further identify genes whose translational efficiencies are frequently controlled by changing usage in the upstream open reading frames during retinal development. These genes are enriched for biological functions highly important to neurons, such as neuron projection organization and microtubule-based protein transport. Surprisingly, we discover hundreds of previously uncharacterized micropeptides, translated from putative long non-coding RNAs and circular RNAs. We validate their protein products *in vitro* and *in vivo* and demonstrate their potentials in regulating retinal development. Together, our study presents a rich and complex landscape of translational regulation and provides novel insights into their roles during retinogenesis.

## INTRODUCTION

The vertebrate retina is a specialized part of the central nervous system with diverse cell types, high-level organization, and an evolutionarily conserved structure (1). It can serve as an ideal model to study neural development, such as deciphering the developmental gene regulatory patterns and understanding mechanisms of morphogenesis formation and specificity (2). To date, genome-wide molecular characterization of retinogenesis has been understood using transcriptomic (3), epigenomic (4), and proteomic (5) approaches and has identified many molecules that play important roles in regulating the development of retina.

The central dogma of molecular biology states two major steps during the detailed residue-by-residue transfer process of genetic information: transcription and translation, by which information encoded in DNA flows into RNA via transcriptional regulation and ultimately to proteins via translational regulation (6). Similar to transcription, translation involves a series of highly temporally orchestrated events directed by cis-elements and trans-factors (7). Increasing evidence emphasizes the importance of translation of gene expression (8,9). Dynamic, tight, and coordinated translational regulation can conduce to the growth of multicellular organisms, particularly during a rapid morphological transition, such as development of red blood cell (10), embryonic stem cell differentiation (11), cortical neurogenesis (12), myogenic differentiation (13), and spermioteleosis (14). In addition, it is suggested that translational regulation may influence plasticity of visual pathway development and function (15). Nevertheless, a systematic, genome-wide analysis of gene translation for retinogenesis is lacking at present.

Recently, accumulating evidence has shown that a fraction of small open reading frames (ORFs) within putative long non-coding RNAs (lncRNAs) are translated to encode functional micropeptides (16). For example, translational product of a lncRNA, *Dworf*, is of critical importance in regulating contraction-relaxation cycles in muscle (17). More recently, some circular RNAs (circRNAs) have been shown to encode bioactive micropeptides, with specific cellular and physiological functions (18,19). These unexpected findings have further emphasized the complex translational regulation in modulating gene expression. However, it is entirely unclear whether the translation of non-coding RNAs and non-canonical ORF-mediated translational control exist during retinal development.

Herein, we conducted the first survey of the translational landscape of neural development in mouse. We applied ribosome profiling (Ribo-seq), mRNA sequencing (mRNA-seq), and circRNA sequencing (circRNA-seq) to the developing mouse retina from the embryonic to the adult stages. We found that translation was dynamically uncoupled with transcription, particularly with larger expression divergence before eye-opening. We revealed diverse regulatory changes fulfilling the requirements of gene expression outputs at different developmental stages. We further detected dynamic changes in translational efficiency (TE) and discovered thousands of upstream ORFs (uORFs), fine-tuning gene translation. Surprisingly, we identified hundreds of actively translated lncRNAs and ribosome-associated circRNAs, which were highly developmental stage specific. We validated their translation *in vivo* and *in vitro* and annotated their potential functions in retinal development. Overall, our study provides a snapshot of complex dynamic patterns of gene translation in mouse retinal development and gains new insights into the regulatory principles of gene translation, particularly non-canonical ORF-mediated translational control, during retinogenesis and neural development.

## MATERIALS AND METHODS

### Animals and tissue collection

Wild-type mice of C57BL/6J genetic background were purchased from Guangdong Medical Experimental Animal Center. Mice from 7 different developmental stages including E13 (Embryonic day), E15, P0 (Postnatal day), P6, P13, P21, and M9 (Month) were euthanized, and the eyes were enucleated immediately after sacrifice, and further retinal tissues from enucleated eyes of each mouse were harvested. In detail, two litters of mice per stage were used to represent two biological replicates. The retinal tissues of mice from the same litter were pooled, snap-frozen in liquid nitrogen, and then stored at −80°C prior to library preparation for next-generation sequencing. Notably, retinal tissues of E13, E15, P0, P6, P13 and P21 were separately used for paired mRNA-seq and ribo-seq; retinal tissues of E15, P0, P6, P21 and M9 were separately used for circRNA-seq; and retinal tissues of M9 were also used for ribo-seq. All experimental procedures were approved by the Animal Ethics Committee of Zhongshan Ophthalmic Center, Sun Yat-sen University (Guangzhou, China; License No: SYXK (YUE) 2018-0189).

### Library preparation and sequencing

Frozen retinal samples per stage were lysed using 1 ml of mammalian lysis buffer (200 µl of 5x Mammalian Polysome Buffer, 100 µl of 10% Triton X-100, 10 µl of DTT (100 mM), 10 µl of DNase I (1 U/µl), 2 µl of cycloheximide (50 mg/ml), 10 µl of 10% NP-40, and 668 µl of nuclease-free water). After incubation for 20 minutes on ice, the lysates were cleared by centrifugation at 10,000xg and 4°C for 3 minutes. The lysate was divided into 300-μl and 100-μl aliquots for ribo-seq and in parallel mRNA-seq, respectively. In detail, for the 300-μl aliquots of clarified lysates, 5 units of ARTseq Nuclease were added to each A260 lysate, and the mixtures were incubated for 45 minutes at room temperature. Nuclease digestion was stopped by additional 15 μl of SUPERase·In RNase Inhibitor (Ambion). Subsequently, the lysates were applied to Sephacryl S400 HR spin columns (GE Healthcare Life Sciences), and ribosome-protected fragments were purified using the Zymo RNA Clean & Concentrator-25 kit (Zymo Research). Ribosomal RNA was depleted using the Ribo-Zero magnetic kit (Epicentre). Sequencing libraries of ribosome protected fragments were generated using the ARTseq™ Ribosome Profiling Kit (Epicentre, RPHMR12126) according to the manufacturer’s instructions. From the 100-μl aliquots of clarified lysates, poly(A)+ RNAs were extracted and purified, and sequencing libraries of poly(A)+ RNAs were then generated using the VAHTSTM mRNA-seq v2 Library Prep Kit from Illumina (Vazyme Biotech, NR601-01) according to the manufacturer’s instructions. The resulting 26 barcoded libraries were pooled and sequenced using an Illumina HiSeq 2500 instrument in the single-end mode and in a randomized order across lanes.

For circRNA sequencing, total RNA was extracted using Trizol reagent (Invitrogen, Carlsbad, CA, USA), the RNA integrity was assessed by Agilent 2100 with RIN number >7.0. Approximately 5 μg of total RNA was used for library preparation, the Ribo-Zero™ rRNA Removal Kit (Illumina, San Diego, USA) was used to deplete ribosomal RNA according to the manufacturer’s instructions, the left RNAs were treated with RNase R (Epicentre Inc, Madison, WI, USA) to remove linear RNAs and to enrich circRNAs. The circRNA sequencing libraries were then generated using TruSeq Stranded Total RNA HT Sample Prep Kit (Illumina RS-122-2203) according to the manufacturer’s instructions with minor modifications. Briefly, cDNAs were reverse transcribed using ProtoScript II Reverse Transcriptase. The ligated products were amplified with PCR by the following conditions: initial denaturation at 95°C for 3 min; 8 cycles of denaturation at 98°C for 15 sec, annealing at 60°C for 15 sec, and extension at 72°C for 30 sec; and then final extension at 72 °C for 5min. At last, the resulting 10 barcoded libraries were sequenced using an Illumina Hiseq X Ten instrument in the paired-end mode.

### mRNA-seq and Ribo-seq alignment

The raw Ribo-seq and mRNA-seq sequencing reads were demultiplexed using CASAVA (version 1.8.2), followed by adapter trimming with Cutadapt (v1.9.1, -e 0.1 -O 6 -m 20) (20) and removal of poor-quality reads with Sickle (v1.33, -q 20 -l 20 -x -n; available at https://github.com/najoshi/sickle). The reads aligned to mouse tRNA and rRNA sequences using Bowtie (v1.0.1, -l 20) (21) were further removed. All remaining reads were mapped to the mouse reference genome (GENCODE, Release M18: GRCm38.p6) using STAR (v2.5.2) with default parameters. Only uniquely mapped reads selected by samtools (v1.6, Phred score >= 20) were used for subsequent analysis.

### Gene quantification and expressed genes definition

Raw counts for different genomic features were obtained using featureCounts from the subread package in Bioconductor (v1.5.0, -t CDS for Ribo-seq data and -t exon for mRNA-seq data) (22). The raw counts from all 24 samples were then combined and normalized together by a pool-based size factor yielded from the DESeq2 R package (23) to minimize the batch effects among samples. After that, the expression level of each gene was estimated as reads per kilobase of transcript per million reads mapped (RPKM) using an in-house R script. At the transcriptome layer, only those genes with RPKM > 1 across two replicate samples of each developmental stage were kept, defined as well-transcribed genes. At the translatome layer, only those genes that are well-transcribed and undergo active translation (see below) were defined as well-translated genes.

### Actively translated ORF detection

Actively translated ORF detection in Ribo-seq data was performed using RiboTISH (v0.2.4) with ‘--framebest’ strategy to select the best candidate ORF (24). To increase statistical power of ORF finder, we merged the aligned BAM files from two replicated samples of each developmental stage using ‘samtools merge’ (v1.6). Upstream ORFs (uORFs) were defined as ORFs originating from the 5’-UTRs of annotated protein-coding genes (that is, with TisType: 5’-UTR); downstream ORFs (dORFs) were defined as ORFs originating from the 3’-UTRs of annotated protein-coding genes (that is, with TisType: 3’-UTR); and lncORFs were defined as ORFs originating from annotated lncRNA genes.

### Genes with dynamical changes detection

Genes with dynamic expression patterns over time were identified by the maSigPro R package, which is specifically designed for the analysis of time-course gene expression data (25). The method is a two-regression step approach where the experimental groups are identified by dummy variables. It first adjusts a global regression model with all the defined variables to identify differentially expressed genes, and in second a variable selection strategy is applied to study differences between groups and to find statistically significant different profiles. Genes with significant temporal changes in retinal development were selected at a FDR < 0.05 and R-squared threshold equal to 0.6.

### Translational efficiency estimation

The TE for each protein-coding gene was estimated as the ratio of the normalized RPKM values of Ribo-seq to mRNA-seq reads in annotated CDS regions (26). Given a high degree of TE correlation between two replicated samples of each developmental stage, particularly when only focusing on those well-transcribed genes (mean Pearson’s correlation coefficient = 0.95), TE values of the two replicates for each protein-coding gene were then averaged to form a final TE for subsequent comparative analysis.

### Differential expression analysis, gene classification, and gene ontology (GO) enrichment analysis

Differential expression analysis was performed using deltaTE (27), which incorporates Ribo-seq and mRNA-seq data. Briefly, deltaTE considers sample-to-sample variance and introduces an interaction term into the statistical model of DESeq2, and the interaction term is used to model condition (that is, developmental stages) and sequencing methodology (that is, Ribo-seq or mRNA-seq). In order to identify significant differences between conditions that are discordant between sequencing methodologies, its generalized linear model incorporates three components: the condition, the sequencing type, and an interaction term containing both. For each gene, a false discovery rate (FDR) threshold of 5% is used to determine statistical significance of the resulting change in RPFs (ΔRPF), mRNA counts (ΔRNA), and TE (ΔTE), respectively, and the ΔRPF, ΔRNA, and ΔTE are combined to determine its regulatory group of belonging. In detail, genes with change in mRNA and RPF levels at the same rate were defined as differentially transcribed genes (DTG), and genes with change in RPF level independent of change in mRNA level, which lead to a change in TE, were defined as differential translational efficiency gene (DTEG). DTGs and DTEGs between adjacent stages could further be categorized into four classes: buffered, intensified, forwarded and exclusive. Specifically, translationally buffered genes have a significant change in TE that offsets the change in RNA; translationally intensified genes have a significant change in TE that bases on the effect of transcription; translationally forwarded genes are DTGs that have a significant change in mRNA and RPF at the same rate and with no significant change in TE. Conversely, translationally exclusive genes are DTEGs that have a significant change in RPF but with no change in mRNA leading to a significant change in TE.

GO enrichment analysis was used to reveal biological functions of differentially expressed genes (DEGs), which was achieved by the ClusterProfiler R package (avaliable at http://bioconductor.org/packages/release/bioc/html/clusterProfiler.html) using all genes as the background. Only those GO terms with false discovery rate (FDR) < 0.01 were regarded as statistically significant.

### Principal component analysis

To define the main contributing layer of gene expression regulation (buffered, exclusive, forwarded, and intensified) for each coregulatory biological function, principal component analysis (PCA) was performed as described in a previous study (19). For each GO term, the relative fractions of four defined classes of differential genes were used as an input for the PCA. The prcomp and fviz_pca_biplot functions from the factoextra R package (avaliable at http://www.sthda.com/english/rpkgs/factoextra) were used for the PCA and visualizing the output of the PCA, respectively. Thus, the placement of each GO term in the PCA plot was based on the directionality of four layers of gene expression regulation.

### Sequence conservation analysis

To calculate evolutionary conservation of non-canonical ORFs in sequence, we downloaded phastCons tracks from UCSC based on 60 vertebrate species (http://hgdownload.soe.ucsc.edu/goldenPath/mm10/phastCons60way, August 2019), and then retrieved the base-by-base conservation scores from “mm10.60way.phastCons60wayPlacental.bw”. For each ORF, the mean PhastCons score at each base was used to evaluate the degree of its cross-species conservation. It varies on a scale from 0 to 1, with 0 indicating to be poor conserved and 1 indicating to be strong conserved.

### CircRNA identification and quantification

Raw circRNA-seq reads were pre-processed with Perl scripts, including the removal of adaptor-polluted reads, low-quality reads (Phred score >= 20) and reads with number of N bases accounting for more than 5%. The clean reads were then mapped to the mouse reference genome (GENCODE, GRCm38.p6) using BWA-MEM (-T 19, v0.7.17; available at http://bio-bwa.sourceforge.net/). Next, two different detection tools were used to identify transcribed circRNAs, namely, CIRI2 (v2.0.6) (28) and CIRCexplorer2 (v2.3.6) (29). To further reduce false positives of circRNA identification, only those circRNAs meeting all of the following three criteria were kept, including (1) having at least 2 unique backsplice junction (BSJ) reads, (2) being simultaneously identified in both tools, and (3) being simultaneously detected in two replicated samples of each developmental stage. Finally, the BSJ reads of CIRI2 were used to quantify the transcriptional level of circRNA using CPM (counts per million mapped reads) (30).

### Identification of Ribosome associated-circRNAs

To determine ribosome-associated circRNAs, we first extracted the 40-base pair (bp) sequences on either side of the backsplice junction site of each transcribed circRNA, and then the sequence was ligated in tandem to generate a pseudo circRNA reference. Next, all Ribo-seq reads that failed to map to the linear reference genome were realigned to the pseudo circRNA reference sequences using Tophat2 (v 2.1.1) with default parameters except N, which was set to 0 (the default is 2) (31). Finally, ribosome associated-circRNAs (ribo-circRNAs) were defined as having (1) at least one unique backsplice junction-spanning Ribo-seq reads and (2) a minimum read-junction overlap of three nucleotides (nt) on either side of the backsplice junction site. Notably, the backsplice junction-spanning Ribo-seq reads might arise from linear, trans-spliced products (trans-spliced RNAs; tsRNAs) rather than circRNA molecules. To further reduce false positive results, we took advantage of our poly(A)+ RNA-seq (mRNA-seq) data to detect BSJ reads and identify potential tsRNAs expressed in each stage, considering that circRNAs are generally non-polyadenylated but tsRNAs are not. The ribo-circRNAs sharing the same splicing junction site with these tsRNAs were then excluded from subsequent analyses.

### CircRNA ORF prediction

To predict putative circRNA-encoded ORF (cORF), the cORF_prediction_pipeline was used with some modifications (32). Briefly, the full-length sequence of each ribo-circRNA was retrieved and multiplied four times to allow for rolling circle translation. All possible cORFs with an AUG-start codon followed by an in-frame stop codon in the exonic sequence were identified and then filtered based on the requirements of a minimum length of 20 amino acids (aa) and of spanning the backsplice junction site. Notably, only the longest cORF was retained for each of the three frames per ribo-circRNA. If a cORF does not contain stop codon, it was defined as an INF (infinite)-cORF, representing that the circRNA could be translated via a rolling circle amplification mechanism.

### Two-dimensional liquid chromatography-tandem mass spectrometry

Retina tissues at E13, E15, P0, P6, P13 and P21 were mixed, fully ground in liquid nitrogen and lysed with lysis buffer L3 (Fitgene Biotech, #FP1801), 0.2% SDS and 1x PMSF (Sangon Biotech, #P0754). Lysates were sonicated on ice and cleared by centrifugation at 12000 rpm for 10 min. The protein extracts were purified by overnight acetone precipitation and re-solubilized using L3, concentration of the samples was determined by Bradford assay. Proteins were reduced using 50 mM TCEP (1 hour at 60°C), alkylated using 55 mM MMTS (45 min at RT in the dark), then the protein sample was loaded on a 10 kDa ultrafiltration tube, washed twice with 8M Urea and three times with 0.25M TEAB. For protein digestion, 50 μl of 0.5M TEAB and trypsin (Promega, enzyme: protein ratio of 1:50) were added to the membrane, reaction was incubated overnight at 37°C. The resulting peptides were first fractionated on a Gemini-NX 3u C18 110Å 150*2.00 mm columns (Phenomenex, #00F-4453-B0) using high-pH reversed-phase chromatography (Dionex Ultimate 3000 RSLCnano) with increasing concentration of acetonitrile, 20 fractions were collected according to the 214 nm absorbance and running time. After vacuum drying, 3 μg of fractionated peptide samples were separated on an Acclaim PepMap RSLC C18 2μm 100Å 75 μm i.d. × 150mm column (Dionex, #160321) and analysed using Thermo Scientific™ Q Exactive™. Full MS spectra from m/z 375-1800 were acquired at a resolution of 70,000 with an automatic gain control (AGC) target value of 3e6 and maximum injection time (IT) of 40 ms. MS/MS spectra were obtained at a 17,500 resolution with an AGC target of 1e5 and maximum injection time (IT) of 60 ms, TopN was set to 20 and NCE/stepped NCE was set to 27. In addition, two unfractionated peptide samples were also analysed by LC-MS/MS using the same parameters as fractionated peptide samples.

### Analysis of mass spectrometry-based proteomic data

Three publicly available proteomic datasets obtained from the PRIDE database (Accession number: PXD003441, PXD002247, and PXD009909) and an in-house MS dataset were used to detect protein products from translatable lncRNAs and circRNAs. The raw data files were analysed using MaxQuant software (v1.6.15.0) (33) against a custom-made database, which combined all mouse sequences from UniProt/Swiss-Prot (MOUSE.2020-08) with sequences derived from translatable lncRNAs and circRNAs, based on the target decoy strategy (Reverse) with the standard search parameters with the following exceptions: (1) the peptide-level FDR was set to 5%, and the protein-level FDR was excluded; (2) the minimal peptide length was set to 7 amino acids; and (3) a maximum of two missed cleavages was allowed. Each search included carbamidomethylation of cysteine as fixed modification methionine oxidation, N-terminal acetylation as variable modifications, but for PXD002247 and our own MS data, a variable modification of deamidation of asparagine and glutamine was also included.

### Functional annotation of lncORF-encoded peptides

#### Conserved domain and protein homology detection

Each of putative micropeptides encoded by lncORFs was annotated against the InterPro database using InterProScan (v5.44) with the default parameters (34). In total, 212 of 603 micropeptides were mapped to known homologous records in the InterPro database, of which 68 were annotated with specific functional domains involving in vital important pathways (**Table S1**).

#### Guilt–By–Association Approach

A total of 198 translatable lncRNAs with dynamical changes in the retinal development were grouped into three clusters by using k-means clustering algorithm. To infer biological functions of each lncRNA cluster, the guilt–by–association approach was used, where (1) Pearson’s correlation coefficient between each lncRNA-mRNA pair was computed in all samples in our dataset; (2) candidate lncRNA-mRNA pairs were selected as those with correlation coefficients > 0.70 and significance level of 0.05 for Pearson’s correlation (FDR<0.05); and (3) protein-coding gene co-expressed with any one lncRNA of each cluster were merged into a union set of genes for GO enrichment analysis.

### Validation of uORF-mediated translation repression

To validate translation repression mediated by uORF, *Neurod1* was selected to construct reporter vector. The 5’-UTR and CDS sequence of 1,242 bp was cloned into pCAG-eGFP, and the main ORF of Neurod1 was Flag-tagged at its C-terminus. The EGFP with an IRES was used as an internal control for transfection and RNA expression. To measure the levels of NEUROD1 protein with a C-terminal FLAG-tag, Neuro-2a cells were transfected with wild-type (WT) or mutant (MUT) reporter plasmids by using Lipofectamine 3000 (Invitrogen). At 24 hours post transfection, cells were passaged to 24 well plate and cultured with complete media (10% FBS) or differentiate media (1% FBS) for 2 days. And then cells were harvested, washed in phosphate-buffered saline (PBS) and lysed in RIPA lysis buffer (Invitrogen) added with Benzonase and protease inhibitor. Proteins were separated by sodium dodecylsulphate-polyacrylamide gel electrophoresis (SDS-PAGE), and then transferred to 0.2 μm polyvinylidene fluoride (PVDF) membranes. The membrane was blocked in 3% nonfat dry milk, and then probed with anti-GFP, anti-FLAG and anti-Histone H3.1 primary antibody. The protein bands were developed with HRP-Substrate ECL (Millipore) detected with the Alliance Q9 system.

### Translation validation of uORFs and lncORFs

#### Plasmid construction

We constructed a series of expression vectors for the detection of non-canonical ORF translation, including 4 uORFs (*u-Rnf10, u-Rnft1, u-Usp8 and u-Zkscan17*) and 10 lncORFs (*Brip1os, Cct6a, Gas5, Malat1, Miat, Peg13, RP23-41oL16*.*1, RP24-112I4*.*1, Six3os1, and Zfas1*), together with two known translatable lncRNAs (*Mrln* and *Dworf*) and one protein-coding gene (*DHFR*) as positive controls. The uORFs and lncORFs (including the predicted 5’- and 3’-UTR) were cloned from mouse retinal cDNAs; *Mrln* were cloned from mouse muscle cDNAs; *Dworf* were cloned from mouse heart cDNAs; and *DHFR* were cloned from Hela cells cDNAs. After obtaining the sequence of these candidates and controls, a HiBiT tag was inserted upstream the stop codon of each predicted ORF by inverse PCR to enable luminescence-based detection of translation products. In order to confirm that peptides were indeed translated from the corresponding ORFs, mutations were introduced in the start codons of predicted ORFs using ClonExpress II One Step Cloning Kit (Vazyme, #C112-02). Additionally, 10 ORFs (*Cct6a, Gas5, Malat1, Mrln, RP23-41oL16*.*1, RP24-112I4*.*1, Six3os1, Zfas1, u-Rnft1 and u-Usp8*) and their mutants were added with 3xFlag tags downstream the HiBiT tag. Sequences of all constructed plasmids were verified by sanger sequencing and the plasmid DNA was extracted using EndoFree Plasmid Midi Kit (CWBIO, #CW2105). The primers and oligos used for cloning, inverse PCR, mutation generation, and epitope tagging were listed in **Table S2**.

#### Luminescence-based detection of translation products

The translation of ORFs were detected using both *in vitro* translation assays (IVT assays) and cultured cells. For IVT assays, TnT^®^ Quick Coupled Transcription/Translation System was used for *in vitro* translation of all HiBiT-tagged ORFs. According to the manufacturer’s instructions, 1 μg of plasmid DNA was used as template for each reaction and the reaction was incubated for 90 min at 30°C. The detection of translated products was performed using Nano-Glo^®^ HiBiT Lytic Detection System (Promega, #N3030) according to the manufacturer’s instructions with minor modifications. In brief, 10 μl IVT products were diluted to 50 μl using nuclease free water, added with 50 μl lytic buffer and mixed by pipet, after 10 minutes incubation at room temperature, the luminescence was measured on a Promega GloMax^®^-Multi Detection System. To further explore the translational potential of the candidates in cellular context, 500ng of plasmid DNA was transfected into N2A, Hela and ARPE19 cells using Lipofectamine 3000. Cells were harvested at 48 hr post-transfection for subsequent analysis. The plate was equilibrated to room temperature, then 300 μl lytic buffer was added to each well and incubated for 10 minutes on orbital plate shaker. The lysates were divided into three tubes and the luminescence of each tube was measured on a Promega GloMax^®^-Multi Detection System.

#### Western blots

The translated peptides were further validated by western blot. Transfected N2A Cells were harvested with 100 μl of RIPA buffer (25 mM Tris•HCl pH 7.6, 150 mM NaCl, 1% NP-40, 1% sodium deoxycholate, 0.1% SDS) added with 1X PIC (Merck, #539131) and Benzonase (NovoProtein, #M046-01B) and incubated 10 min on ice. For detection of peptides from *Brip1os, Miat, u-Rnf10*, and *DHFR*, lysates were added with 5×SDS-PAGE Sample Buffer (GenStar, #E153) and denatured at 95°C for 5 minutes, and samples were loaded on 4–20% Mini-PROTEAN^®^ TGX™ Precast Protein Gel and transferred to a 0.2 μm NC membrane. For detection of the remaining peptides, lysates were added with 2x Novex Tricine SDS Sample Buffer (Invitrogen, #LC1676) and denatured at 85°C for 2 minutes, and samples were loaded on 16.5% GLASS Gel^®^ Tricine gel (WSHT, #TCH2001-16.5T) and transferred to a 0.1μm NC membrane. The blot was carried out using Nano-Glo^®^ HiBiT Blotting System (Promega, #N4210) according to the manufacturer’s instructions.

### Validation of circRNA expression

#### Divergent PCR validation

Divergent primers were designed to amplify the circRNA backsplice junction sequence and retinae cDNA were used as template for divergent PCR (**Table S2**). Divergent PCR was performed using green Taq mix (Vazyme), and the reaction was carried out for 3min at 95°C and 30 cycles of 15 sec at 95°C, 15 sec at 60°C and 30 sec at 72°C. The PCR products were then analysed on 1.5% agarose gels in 1x TAE buffer. All of the PCR products were sanger sequenced with forward and reverse primers to find the backsplice junction sequence.

#### RNase R treatment assay

RNase R treatment and circRNA quantification was performed according to a published protocol (Panda and Gorospe, 2018) with minor modification. In brief, RNA was treated with 20 μl RNase R digestion reaction (2 μg RNA, 1 μl RNase R (Lucigen, #RNR07250), 2 μl 10x RNase R reaction buffer) and control reaction without RNase R. The reactions were incubated for 30 min at 37°C and immediately purified using ZYMO RNA Clean & Concentrator (ZYMO RESEARCH, D7011). The purified RNA samples were eluted in 20 μl of nuclease free water, and 12 μl of RNase R treated RNA and control RNA were used for reverse transcription. Quantification of circRNAs were performed using iTaq Universal SYBR Green Supermix (BioRad, #1725124) according to the manufacturer’s instructions. The qPCR reactions were prepared as follows: 0.1 μl cDNA, 10 μl of SYBR Green Supermix, 1.2 μl primer mix (5 μM each) and 6.8 μl nuclease-free water. Reactions were carried on CFX Connect Real-Time PCR Detection System for 2 min at 95°C and 40 cycles of 5 sec at 95°C and 20 sec at 60°C followed by melting curve analysis. The enrichment of RNA after RNase R treatment was calculated using the delta CT method, mouse gene Gapdh and Rps14 were used as linear control.

## RESULTS

### Ribo-seq, mRNA-seq, and circRNA-seq on the mouse retinal tissue

To understand the translational regulation of gene expression during mouse retinogenesis, we performed Ribo-seq and mRNA-seq to generate translatome and transcriptome profiles of mouse retina at six developmental stages, including E13, E15, P0, P6, P13, and P21, temporally spanning two major developmental events in retina, birth and eye opening (day11-12) (**Fig. 1A**). We also performed circRNA-seq specifically for transcriptomic profiling of circRNAs at E15, P0, P6, P21, and M9. All the sequencing experiments were done with two biological replicates. In total, the Ribo-seq, mRNA-seq, and circRNA-seq yielded 1.17, 0.37, and 0.87 billion raw reads, with an average of around 83.25, 30.98, and 87.38 million reads per library, respectively (**Table S3**).

**Figure 1.**
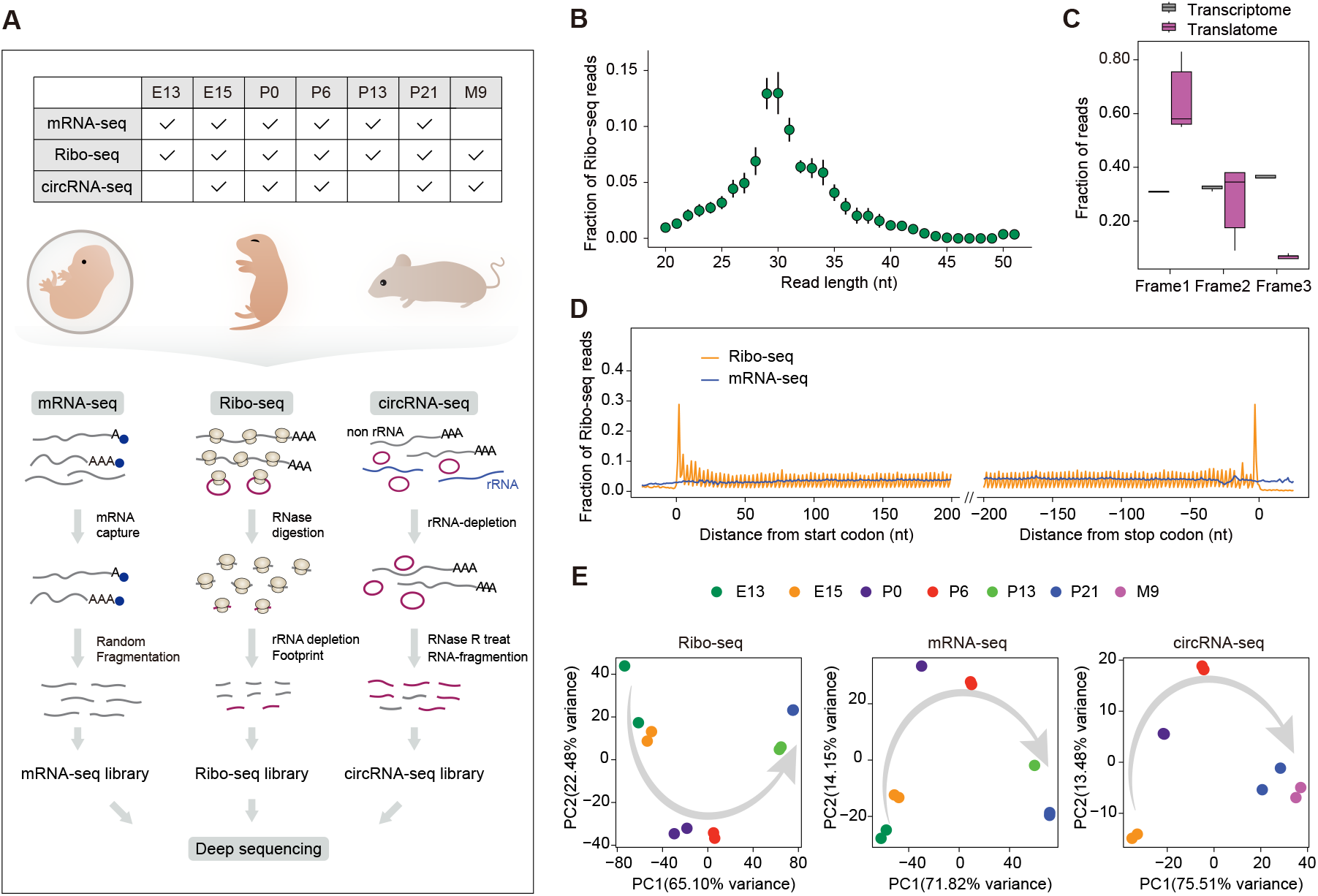
Overview of gene expression of the developing mouse retina. **(A)** A schematic illustration of the experimental design. **(B)** Length distribution of RPFs that mapped to CDSs of protein-coding genes, with a peak at 29-30 nt. **(C)** Frame distribution of Ribo-seq and mRNA-seq reads among all the CDSs, showing a clear frame preference for Ribo-seq reads and a uniform frame distribution for mRNA-seq reads. **(D)** Metagene analysis of read distribution around the start and stop codons of the CDSs, showing a 3-nt periodicity of Ribo-seq reads. **(E)** Principal component analysis of Ribo-seq, mRNA-seq, and circRNA-seq data sets, respectively.

Ribosome-protected fragments (RPFs) generated from the Ribo-seq had a typical length range of 25- to 35-nucleotide (nt), tightly distributed around a peak of 29- to 30-nt (**Fig. 1B** and **Fig. S1A**), a preference mapped to annotated coding sequences (CDS) and 5’ untranslated region (**Fig. S1B**), a strong bias toward the translated frame (**Fig. 1C** and **Fig. S1C**), and a characteristic three-nucleotide (3-nt) periodic subcodon pattern (**Fig. 1D** and **Fig. S1D**). As expected, these characteristics were absent in the mRNA-seq datasets. There was a high degree of agreement between biological replicates, with an average Pearson’s correlation coefficient of 0.981, 0.993 and 0.990 for Ribo-seq, mRNA-seq and circRNA-seq, respectively. Principal component analysis (PCA) showed that the samples had a clear segregation of development stages (**Fig. 1E**) and the samples from the same stage were more similar to each other than the ones from the different stages (**Fig. S1G**). In addition, expression of some known marker genes in retinal development on our datasets, such as *Otx2, Pax6*, and *Neurod4*, well agreed with previous studies (**Fig. S2**) (35-37). Collectively, these results demonstrated that our experimental data were of high quality.

### Coordination of translational and transcriptional regulation during retinal development

Based on the Ribo-seq and mRNA-seq datasets, we identified an average of 11,150 well-translated protein-coding genes and 12,458 well-transcribed protein-coding genes per stage (**Fig. 2A**, see **Methods**). Most of the well-translated and well-transcribed protein-coding genes were shared across all the stages (**Fig. 2B-C**). The shared genes underwent dynamic expression changes during retinal development, with 91.4% (9,294 of 10,172 at the transcriptional level; **Fig. S3A**) and 76.3% (6,961 of 9,119 at the translational level; **Fig. S3B**) genes showing significant changes in temporal differential expression (FDR< 0.05; see **Methods**), including many well-known transcription factors directing retinogenesis such as *Rax, Crx, Otx2, Vsx2*, and *Neurod1* (**Fig. S3C**). Correlation analysis showed that transcriptome and translatome were more different at the early stage but became more similar after eye-opening, suggesting uncoupled changes between transcriptome and translatome during retinal development (**Fig. 2D**).

**Figure 2.**
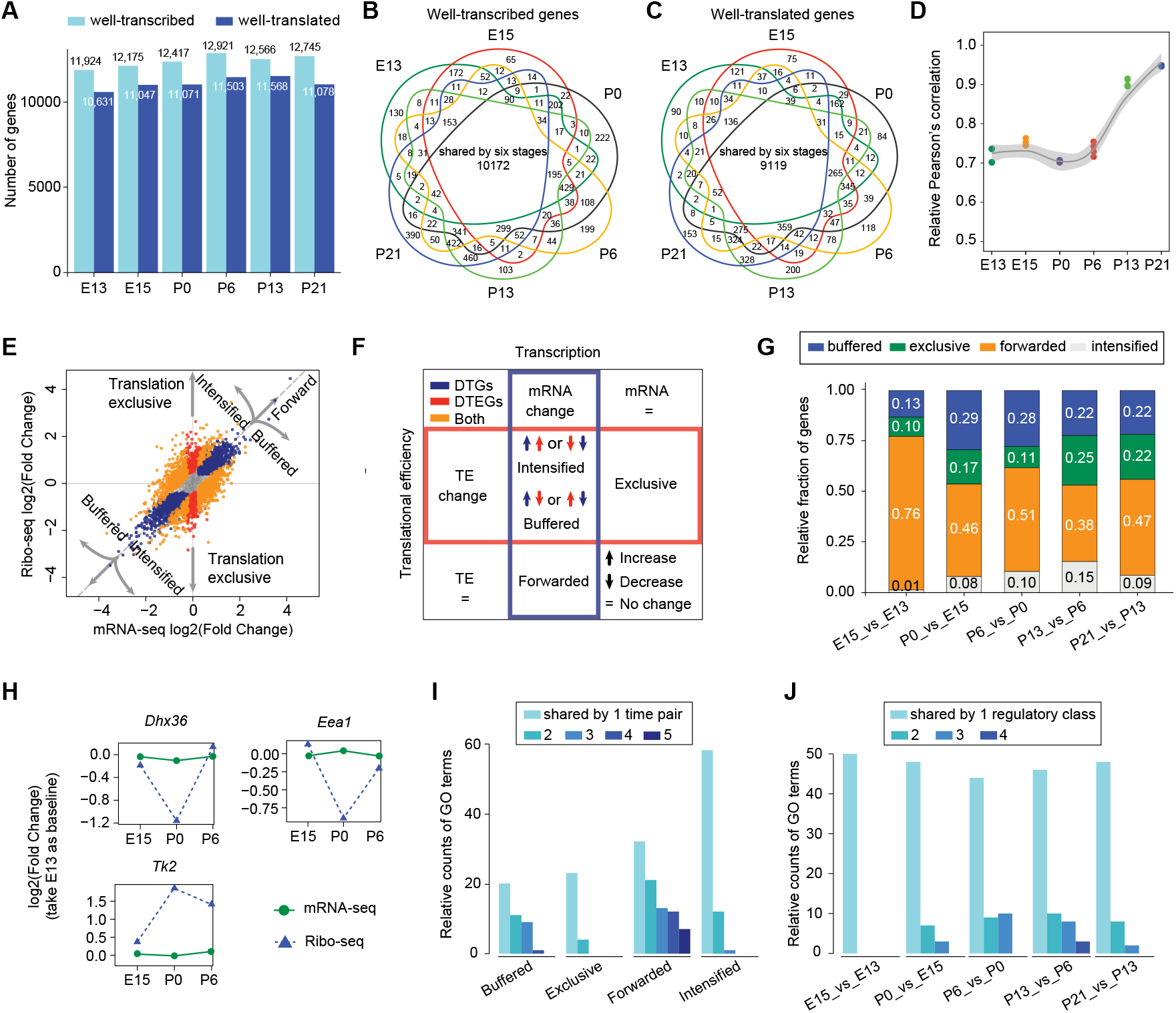
Transcriptional and translational characterization. **(A)** Number of well-transcribed and well-translated protein-coding genes in each stage. Overlap of **(B)** well-transcribed and **(C)** well-translated genes across different developmental stages. **(D)** Correlation of expression levels between transcriptomes and translatomes throughout development. **(E)** Log-fold changes in the mRNA and ribosome occupancy, taking E15 versus P0 as an example. **(F)** The interplay between DTGs and DTEGs showing categories of gene expression regulation. **(G)** Relative fractions of four categories of genes in each pairwise comparison between adjacent stages. **(H)** Ribosome occupancy and mRNA changes of three well-studied genes: *Dhx36, Eea1* and *Tk2* between E13 and P6. Lines display fold changes at Ribo-seq (blue) and mRNA-seq (green) level based on E13. **(I)** Overlap of enriched GO terms for the same category of genes across all five comparisons of adjacent stages. Colours represent degrees of term overlap among the five comparisons of adjacent stages (that is, E15_vs_E13, P0_vs_E15, P6_vs_P0, P13_vs_P6, and P21_vs_P13). **(J)** Overlap of enriched GO terms for different categories of genes in each adjacent stage comparison. Colours represent degrees of term overlap among the four regulatory types of gene expression: buffered, exclusive, forwarded and intensified.

To capture the overall transcriptional and translational changes in gene expression, we integrated mRNA-seq and Ribo-seq datasets to perform differential expression analysis between adjacent stages using deltaTE (**Fig. 2E**, FDR < 0.05; see **Methods**). We observed that the total number of differentially expressed genes (DEGs) gradually increased with development until a peak between P6 and P13, up to a total of 5,753, then followed by a dramatic decline. This pattern was consistent with the progression of retinal differentiation and maturation (**Fig. S3D**). To capture the specific translational changes independent of changes in transcription, we further identified differential translational efficiency genes (DTEGs) and combined the differential direction of these DEGs and DTEGs to categorize them into four distinct regulatory classes: forwarded, exclusive, buffered, and intensified (**Fig. 2F** and **Table S4**; see **Methods**). The forwarded genes have RPF changes that are explained by the mRNA changes. The exclusive genes have changes in TE without mRNA changes. The buffered genes have changes in TE that offsets the mRNA changes and the intensified genes also have changes in TE that amplifies the mRNA changes.

We observed that differences in transcription were not always forwarded to the translational level, and on average, more than 38% of differentially transcribed genes were translationally buffered or intensified (**Fig. 2G**). Of these, translational buffering was more prominent than translational intensification, which further emphasized the existence of extensive translational regulation that shaped gene expression changes during retinal development. Moreover, translational regulation could also influence gene expression independently, with an average of 517 differential genes found in each comparison of adjacent stages whose changes occurred exclusively at the translational level without underlying mRNA changes. For instance, between E15 and P6, ribosomal occupancy significantly changed for many well-studied genes related to the functionality of neurons, such as *Dhx36* that helps specific microRNA localize to neuronal dendrite (38), *Eea1* that restores homeostatic synaptic plasticity (39), and *Tk2* that ensures neuronal function *in vivo* (40) (**Fig. 2H**).

Notably, genes in the same regulatory class exhibited developmental stage-specific functional enrichment (**Fig. 2I**), while genes in the different regulatory classes exhibited class-specific functional enrichment (**Fig. 2J**). For instance, “neuron projection organization” which is critical for the establishment of retinal visual function was particularly enriched for the exclusive genes during P6 to P13 and intensified genes during P13 to P21. Overall, these results revealed that both translational and transcriptional regulation had important but different roles in the development of retina.

### Functional importance of translational regulation

We next attempted to examine the relative contribution of translational and transcriptional regulation for the biological functions important to retinogenesis. We first performed GO enrichment analysis for the above-detected differentially transcribed and translated genes (FDR < 0.01, **Table S5**; see **Methods**) and identified specific functions enriched throughout the development of the mouse retina (**Fig. 3A**). Particularly, many of these functions were associated with synapse formation and synaptic transmission at neuromuscular junctions, such as “visual system development”, “postsynapse assembly”, “postsynapse organization”, and “signal release from synapse”. We also observed that some biological functions showed stage-specific enrichment. For instance, “positive regulation of chromosome separation” and “microtubule-based protein transport” were exclusively enriched between P0 and P6 as well as between P6 and P13, respectively.

**Figure 3.**
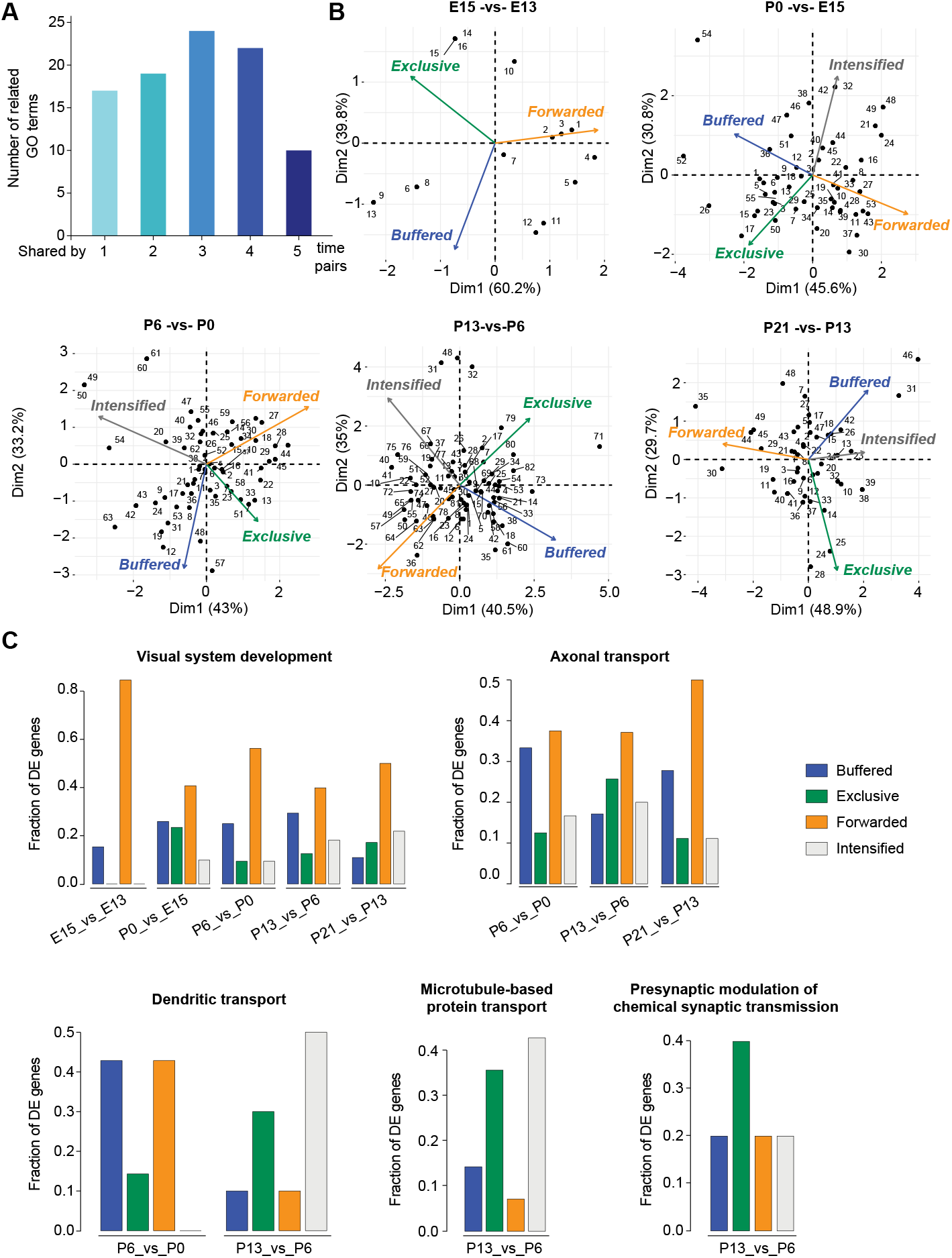
Dissecting Translational regulation in the developing mouse retina. **(A)** Frequency distribution of occurrences of enriched GO terms in all comparisons of adjacent stages. Colours represent overlap degrees of term among the five comparisons of adjacent stages (that is, E15_vs_E13, P0_vs_E15, P6_vs_P0, P13_vs_P6, and P21_vs_P13). **(B)** Scatter plot of PCA of each developmental stage separating the manifestations of individual GO terms within the global regulatory programs (see **Methods**). Each numbered point represents a functional GO term, and its position along each axis indicates the relative contribution of transcriptional and translational regulation to the overall differential patterns. For each arrangement, the assigned GO term can be found in **Table S5. (C)** Examples of stage-specific coregulatory functional arrangements. The barplots show the relative fractions of expressed genes in retina with distinct regulatory modes.

We then calculated the relative percentages of regulatory classes (that is, forwarded, exclusive, intensified, and buffered) of differential genes for each functional term and performed a PCA to dissect their contributions to the overall expression change of each term. The manifestations showed that these important biological functions were under different degrees of translational and transcriptional regulation (**Fig. 3B**). For instance, the “visual system development” was under combinatorial regulation of transcriptional and translational control mechanisms. At the early stage, this function was subjected to the major transcriptional forwarded regulation, but as development progressed, transcriptional regulation gradually weakened and translational regulation, particularly translationally exclusive and intensified regulation, gradually strengthened (**Fig. 3C**). Signal transmission-related function “axonal transport”, enriched frequently after birth, was mainly subjected to translational regulation. Specifically, about 62.7% of total contributions in “axonal transport” could be attributed to translational regulation (exclusive + intensified + buffered), of which 72.9% came from exclusive and intensified regulation. Besides these, the stage-specific enriched functions that played important roles in the homeostatic maintenance of mature synapses, such as “microtubule-based transport” and “presynaptic modulation of chemical synaptic transmission”, were mainly under translational intensified and exclusive regulation, respectively. Our results showed a rich and complex regulation of gene expression during retinal development.

### Dynamics of translational efficiency and contribution of regulatory uORFs

Due to long half-lives (>2 h) for the majority of eukaryotic mRNAs, regulation of their encoded proteins is achieved by controlling mRNA TEs and protein degradation rates (41). We detected a total of 5,945 differential translational efficiency (DTE) genes between adjacent stages (see **Methods**). Unsupervised clustering by *k*-means revealed the temporal dynamics of TE that were categorized into seven clusters (**Fig. 4A**). Each cluster was composed of distinct genes with specific biological functions (**Table S6**). Particularly, the 1049 DTE genes in the cluster C were significantly enriched in “synapse organization”, “postsynapse organization”, and “neuron projection organization”. TEs of these genes in the cluster C were specifically enhanced around E15 (embryonic wave) and P13 (postnatal wave), which would help promote neuronal differentiation, thereby facilitating production of functionally active neurons. The 925 DTE genes in the cluster E were significantly enriched in “mRNA/tRNA 5’-end processing”, “midbody abscission”, and “microtubule-based transport”. Notably, these genes showed peak TE at P6 and might play a critical role in maintaining retina homeostasis and later neurogenic divisions, given typical characteristics of extensive alternative splicing (42) and active cell division events (43) during this period of retinal development. In addition, the 816 DTE genes in the cluster B showed peak TE at the embryonic period and were mainly enriched in “mitotic nuclear division”, “negative regulation of chromatin organization”, and “positive regulation of chromatin organization”, fulfilling the requirements for early neurogenic divisions.

**Figure 4.**
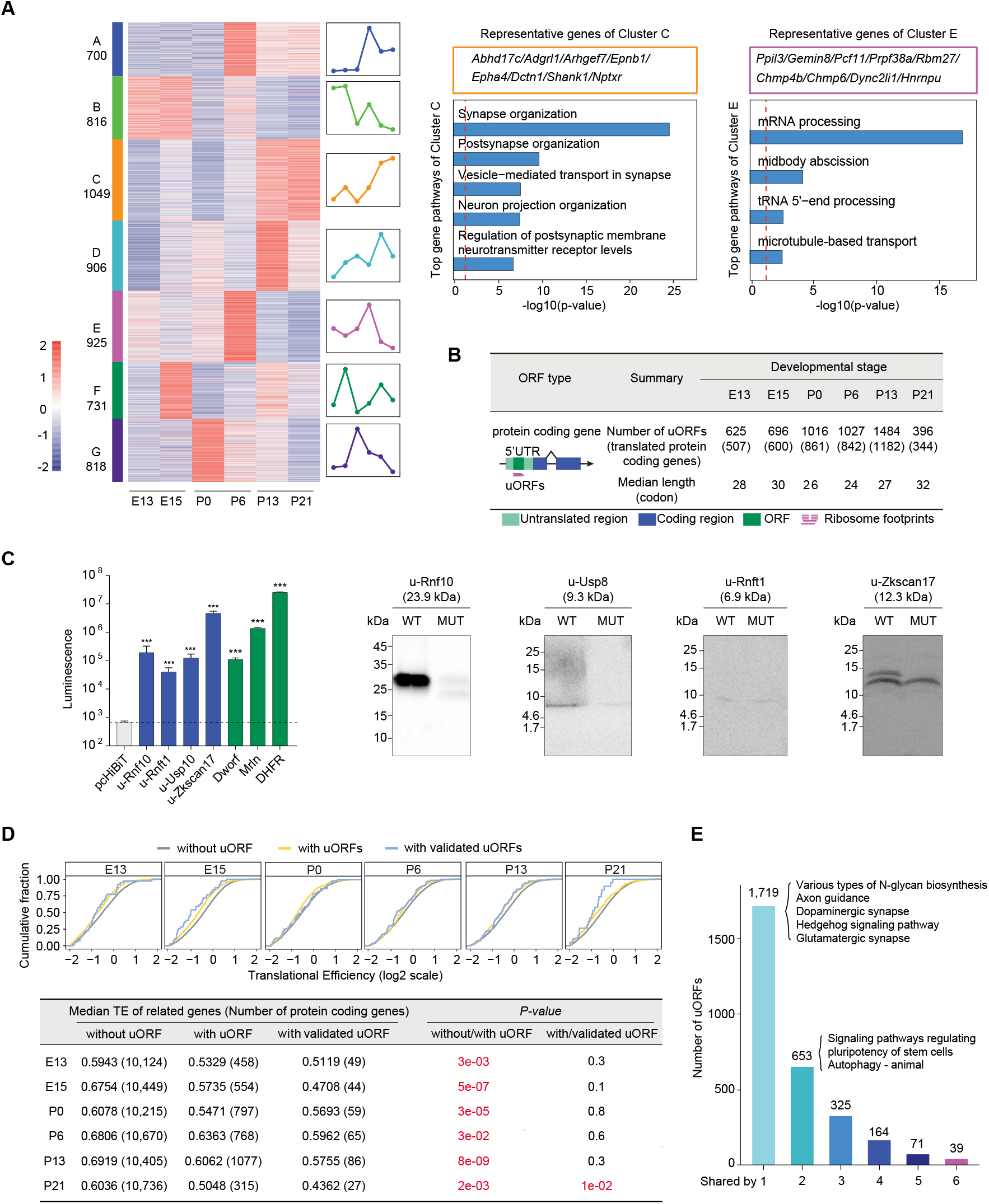
Analysis of translational efficiencies across retinal development. **(A)** Hierarchically-clustered heatmap of TE of 5,945 protein-coding genes (FDR < 0.05; as described in (27)). Different colours indicate different clusters; numbers indicate significant DTE genes in the cluster; and temporal patterns of TE changes are portrayed through lines. Representative genes and enriched GO terms in the two largest clusters are shown on the right. **(B)** Number of actively translated uORFs detected in each stage by using Ribo-TISH. The uORF distribution is shown below. **(C)** Translation validation of uORFs. Left: luminescence of uORFs in IVT assays, where all candidates were compared with the negative control and the dashed line indicates the luminescence of the negative control (negative control: pcHiBiT, positive controls: Dworf and Mrln, protein-coding control DHFR; ns, P > 0.05; *, P < 0.05; **, P < 0.01; ***, P < 0.001); Right: western blots of uORF coded peptides (HiBiT tagged) and corresponding ATG mutants. **(D)** Left: Comparison of TEs between protein-coding genes without, and with and with validated uORFs in each stage; Right: summary of TE between genes without/with/with validated uORF (Wilcoxon rank-sum test). **(E)** Frequency distribution of occurrences of uORFs during retinal development, showing developmental stage-specific usage. Colours represent numbers of uORFs with different usage frequencies during mouse retinal development.

Previous studies have shown that regulatory elements located on mRNA transcripts might affect gene translation, particularly uORFs in the 5’-UTRs (44-47). In total, we detected 2,971 actively translated uORFs in 2,123 protein-coding genes (**Fig. 4B** and **Table S7**; see **Methods**). We randomly selected four uORFs and successfully validated their translated products by IVT assays (**Fig. 4C**). In further support of uORF translation, we used mass spectrometry (MS)-based proteomics data to provide direct *in vivo* evidence for translation of 181 uORFs (**Table S7**; see **Methods**). Notably, only a minority of uORFs (327) showed strong evidence of conservation (PhastCons score>0.95; see **Methods**). This result indicated the majority of uORFs lacking signs of selective pressure to maintain their amino acid sequences, consistent with the findings of previous study (44), suggesting that uORF function might be largely independent of their translated products.

We next sought to understand the effect of uORFs on CDS translation. By comparing the CDS TEs of genes with and without uORFs, we consistently observed a significant uORF-mediated translational repression (**Fig. 4D**, Wilcoxon rank test). One representative uORF residing in *Neurod1* was experimentally validated to further confirm the repressive effect of uORFs on downstream CDS translation (**Fig. S4A**). Comparative analysis of the CDS TEs of genes with single versus multiple uORFs further showed multiple uORFs tended to have an additive repressive effect (**Fig. S4B**), with similar trends observed in five out of six developmental stages displaying significant non-randomness (Fisher’s exact test, *P*-value = 0.0313). Given that the majority (57.9%, 1,719/2,971) of uORFs were found exclusively in a single stage, suggesting that uORF-mediated translational regulation could occur in a stage-specific manner (**Fig. 4E**). The GO enrichment analysis revealed that uORF-containing genes participated in many important biological processes required for retinal development, such as “axon guidance”, “dopaminergic synapse”, and “signaling pathways regulating pluripotency of stem cells” (**Fig. 4E** and **Table S8**). Moreover, we detected actively translated dORFs in the 3’-UTRs. In addition to thousands of translated uORFs, however, a total of only 266 dORFs were detected in 217 protein-coding genes, with an average of 64 dORFs per stage. Given that translation of dORFs significantly enhance translation of their corresponding canonical ORFs (48), we compared TEs of genes with and without dORFs, but did not observe significant enhancive effect of dORFs on their CDS translation possibly due to their much smaller number per stage (**Fig. S4C**).

### Translation of long noncoding RNA genes

Micropeptides encoded by presumed long noncoding RNAs (lncRNAs) have frequently been overlooked, and their prevalence and potential function in retinal development, even in neural development, remain unknown. To discover translated lncRNAs in the developing mouse retina, we searched for actively translated ORFs in lncRNAs (lncORFs) (see **Methods**). In total, we identified 603 unique lncORFs in 290 lncRNAs, with a median length of 48 amino acids (aa) per lncORF (**Fig. 5A** and **Table S9**). Majority of these lncORFs (57.5%, 347/603) could be found in another translatome dataset of mouse retina (GSE94982), including previously well-characterized micropeptides *B230354K17Rik* (49) and *Crnde* (50). Comparing with MS-based proteomics data, we provided direct *in vivo* evidence for translation of 75 out of the 290 lncRNAs (**Fig. S5A** and **Table S9**; see **Methods**).

**Figure 5.**
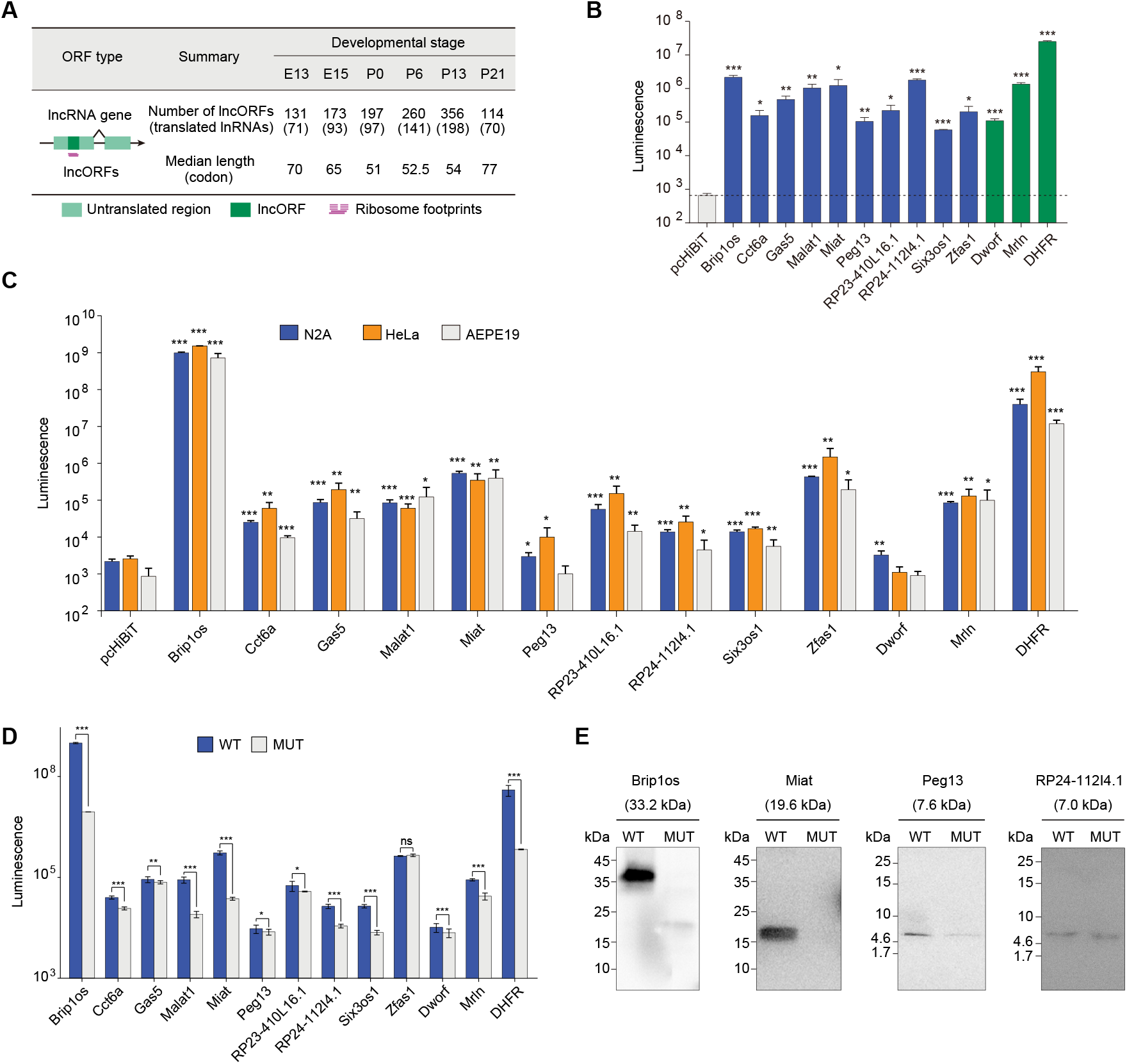
Translation of lncRNAs. **(A)** Number of actively translated lncORFs detected in each stage. **(B)** Luminescence of 10 candidate lncORFs and controls in IVT assays. All candidates are compared with the negative control and the dashed line indicates the luminescence of the negative control. **(C)** Luminescence of 10 candidate lncORFs in N2A, Hela and ARPE19 cells. All candidates are compared with the negative control. **(D)** Luminescence of all lncORFs and corresponding ATG mutant ORFs in N2A cells. **(E)** Western blots of lncORF coded peptides (HiBiT tagged) and corresponding ATG mutants. Note: negative control: pcHiBiT, positive controls: *Dworf* and *Mrln*, protein-coding control *DHFR*; ns, *P* > 0.05; *, *P* < 0.05; **, *P* < 0.01; ***, *P* < 0.001.

To further experimentally validate the translation products, we first performed IVT assays for 10 randomly chosen lncORFs. Considering the relatively low molecular weight of potential micropeptides, we specifically fused a 11-amino-acid HiBiT epitope tag to the C-terminal of each lncORF that could produce bright and quantitative luminescence through high affinity complementation with LgBiT (51). The quantifiable results demonstrated that all of them successfully produced micropeptides in IVT assays (**Fig. 5B**). Furthermore, we separately transfected the expression vectors of these lncORFs into N2a (Neuro-2a), Hela, and ARPE19 cells and detected micropeptide products of all of them in at least two cell types (**Fig. 5C**). Subsequent start codon mutation of these lncORFs prevented their translation or caused truncated translation, as evidenced by significant decreases in the luminescence intensity (**Fig. 5D** and **Fig. S5B**). Loss of signal in the predicted size range of micropeptide products was additionally confirmed by western blot analysis (**Fig. 5E** and **Fig. S5C**).

Translation of lncRNAs showed a strong stage-specificity, with an average of 112 lncRNAs per stage detected to undergo active translation but only 19 shared across all developmental stages (**Fig. S5D**). Among the 290 translated lncRNAs, 198 exhibited significant temporal expression changes, of which 12 were found exclusively in translatome, suggesting that lncRNA translation might independently contribute to changes in developmental programs (**Fig. S5E-F**). We also sought to understand the tissue-specificity of these 290 translated lncRNAs. We conducted Ribo-seq and mRNA-seq for six tissues in mouse and found 172 (59.3%), 102 (35.2%), 133 (45.9%), 79 (27.2%), and 128 (44.1%) of them that also had translation evidence in the brain, heart, kidney, liver, and lung, respectively, with 18 translated in all six tissue types (GSE94982). Importantly, the majority of these lncRNAs (86.1%-91.5%) shared at least one identical lncORF with those detected in the retina. These results suggested that some of the translated lncRNAs detected in the developing retina had potential multi-tissue activities. Functional annotation of these micropeptides translated from lncRNAs using InterProScan 5 revealed 68 (∼11.3%) of 603 lncORFs with identifiable features found in known proteins (**Table S1**), such as conserved domains, protein families, and functional sites, some of which even could be assigned to defined molecular functions. For instance, micropeptide encoded by *Ptpmt1* participated in “protein tyrosine/serine/threonine phosphatase activity” and micropeptide encoded by *RP23-95L9*.*6* participated in “nucleic acid binding”. Our analysis suggested that these lncORFs had functional relevance which could be tested in the future study.

### Translation of circRNAs

An increasing number of studies have indicated that circRNAs can also be translated into detectable peptides with physiological functions (52). We used circRNA-seq and Ribo-seq data to explore translation potentials of circRNAs. Using stringent identification and filtering strategies (**Fig. S6A**; see **Methods**), 255 of 28,910 transcribed circRNAs with at least two supporting back-spliced footprints were defined as ribosome-associated circRNAs (ribo-circRNAs) (**Table S10**). Notably, the majority (82.0%) of them were found in the stages after eye opening, suggesting their potential functional importance for the later stage of retinal development (**Fig. 6A**).

**Figure 6.**
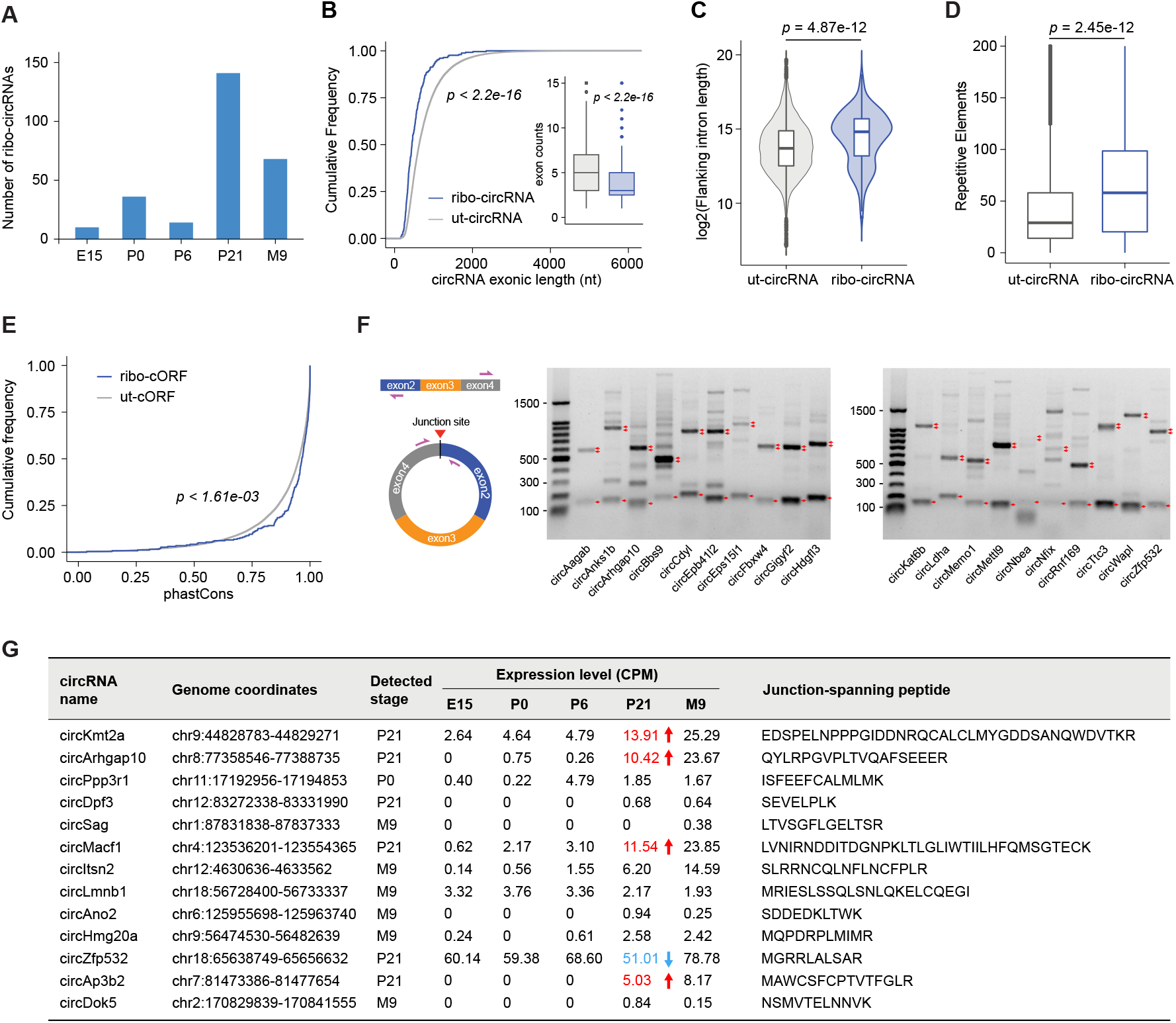
Translation of circRNAs. **(A)** Number of ribo-circRNAs identified in each stage. **(B)** Cumulative plot of exonic length for untranslated circRNAs (ut-circRNAs, black) and ribo-circRNAs (green). The inserted box plot showing exon number differences between ut-circRNAs and Ribo-circRNAs. **(C)** Violin plot of flanking intron length for ribo-circRNAs and ut-circRNAs. **(D)** Box plot comparing the repetitive elements in the flanking introns between ribo-circRNAs and ut-circRNAs. **(E)** Cumulative plot of sequence conservation (phastCons score) for ribo-cORFs (green) and ut-cORFs (black). **(F)** Schematic overview of circRNA divergent primer design (left) and experimental validation of the rolling circle cDNA products from circRNAs (right). **(G)** Summary of the 13 ribo-circRNAs producing putative micropeptides. The stages that circRNAs differentially transcribed were indicated by arrows, red represents up-regulated and blue represents down-regulated.

Ribo-circRNAs had distinguishing properties from untranslated circRNAs (ut-circRNAs). Ribo-circRNAs had significantly shorter exonic length and fewer exons than the ut-circRNAs, with at least 29% and 40% decreases in the median length and number of exons, respectively (**Fig. 6B**; Wilcoxon rank-sum test, *P*-value < 2.2e-16), in agreement with previous observations in human cell lines (53). Ribo-circRNAs had significantly longer flanking introns (**Fig. 6C**; Wilcoxon rank-sum test, *P*-value = 4.87e-12) that harboured more repetitive elements than ut-circRNAs (**Fig. 6D**; Wilcoxon rank-sum test, *P*-value = 2.45e-12). In addition, evolutionary conservation of micropeptides translated from ribo-circRNAs was significantly higher than that of ut-circRNAs (**Fig. 6E**; Wilcoxon rank-sum test, *P*-value < 1.61e-03), suggesting that they were likely to have functional roles, given that strong conservation in sequence is a general indicator of important biological function (54).

Next, we randomly selected 20 candidates including 19 ribo-circRNAs and 1 ut-circRNA and performed Sanger sequencing of RT-PCR products using divergent primers to confirm the back-splice junction sites of these circRNAs (**Fig. 6F** and **Fig. S6B**). After RNase R treatment, all these circRNAs were resistant to digestion with RNase R exonuclease, validating the existence of these ribo-circRNAs (**Fig. S6C**). Using MS-based proteomics data, we further provided direct *in vivo* evidence for translation of 13 ORFs spanning the splice junctions. After minimizing the effect of trans-spliced RNAs (see **Methods**), these micropeptides were more likely to derive from ribo-circRNA themselves. Of the 13 ribo-circRNAs producing putative micropeptides, five were found to be differentially transcribed (**Figure 6G**), including *circKmt2a, circArhgap10, circMacf1, circAp3b2* (up-regulated in P21 compared to P6), and *circZfp532* (down-regulated in P21 compared to P6), emphasizing that their translation might be important for retinal development.

## DISCUSSION

Our integrative analysis of transcriptome and translatome during retinal development revealed specific changes in translation and regulatory roles of translational regulation in retinogenesis. We found that retinogenesis is accompanied by dynamic, rapid and coordinated changes in gene translation and translational regulation. Thousands of important regulatory protein-coding genes were subjected to significant changes in the translational level in a stage-specific manner, potentially redefining functional architecture and diversity of the retinal cells.

Specific translational regulation could be achieved by controlling TEs, thereby determining quantitative differences in protein abundance during retinogenesis. TE dynamics during developmental transition could further be triggered by some regulatory elements, such as the most commonly used uORFs. Within 5’-UTRs, uORF-mediated translational control was a vitally regulatory mechanism for gene expression in mammals (55). Our findings revealed that uORFs provided functionally important repression for many key genes associated with retinal development, such as an uORF residing in *Otx2* gene only at P0 and two uORFs residing in *Nrl* gene at E13, P0, P6, and P13, displaying stage-specific influence on retinogenesis. This repressive effect of uORF on translation of primary CDS appeared to occur in a dose-dependent manner. Notably, many properties may contribute to an uORF’s role in translational regulation, including the length of the 5’UTR, the secondary structure and GC content, as well as the strength of the surrounding Kozak context, the uORF length, and conservation, which have been substantially discussed in detail in previous studies (46,56,57). A uORF with a suboptimal context may benefit leaky scanning and hence allow its main CDS translation (58). Due to these complexities of uORF-mediated translational regulation, this might partially explain why the change of translation efficiency by uORF is not significant at certain stages. Moreover, many uORF-encoded peptides were here detected. Recent studies have suggested that they may modulate translation or have other cellular functions. For instance, uORF-encoded micropeptides act *in cis* to cause ribosome stalling or limit ribosomal access to the main CDS or act *in trans* to suppress translation (59). uORF-encoded peptides can serve as MHC class I ligands, thereby contributing to the antigen repertoire and possible immunogenicity (60). Detailed physiological functions and underlying mechanisms of uORFs in shaping retinogenesis via producing micropeptides remain to be elucidated.

Surprisingly, we additionally discovered pervasive translation of lncORFs from lncRNAs. We experimentally confirmed their translation products and showed that these micropeptides had regulatory potential and biological relevance. For instance, we found that the micropeptide encoded by *RP23-15O6*.*3* contained C2H2-type zinc fingers. The micropeptide encoded by *AC166710*.*3* shared a common structure with ribosomal L18e/L15P superfamily. Functional enrichment analysis revealed potential roles of these translated lncRNAs in retinal development (**Fig.S5F**). One representative lncRNA was *Miat* (also known as *Rncr2* or *Gomafu*) that contributed to mitosis of retina progenitor cells, illustrating its essential role for retinogenesis (61). Besides the direct involvement of retinal development, some of the encoded micropeptides may serve as regulatory elements to play important roles in modulating neighbouring gene activity. This is partially supported by the observation that neighbouring genes of many translated lncRNAs such as *Six3os1* (IVT assays and *in vivo* translation), *Vax2os, Otx2os1, Pax6os1*, and *Zeb2os1* (MS detection) were important determinations of retinal cell fate (62). In our study, micropeptides were only evident for a relatively small subset of uORFs, dORFs, and lncORFs. The possible reasons behind this were: (1) the majority of their encoded peptides might lack signs of selective pressure to maintain their amino acid sequence; (2) their encoded peptides may be relatively short lived and rapidly degraded; (3) their encoded peptides might exist at only very low levels within the cell; and (4) our high-pH reversed-phase fractionated retinal proteomics data did not cover the full peptide isoelectric point range so that it was unable to explore the entire tryptic peptidome. To obtain a comprehensive map of hidden micropeptides during retinal development, innovative technologies are needed.

Notably, 13 micropeptides encoded by non-canonical ORFs spanning the splice junctions were detected using MS-based proteomics data. These micropeptides might be partially associated with the functions of their host proteins due to the existence of overlapping sequences between them, likely competing with (63) or protecting (64) their host proteins. It should be noted here that circRNAs possess coding potential and can indeed be translated, but translation may originate from their linear trans-spliced RNA by-products (tsRNAs) (65). Although we take advantage of our poly(A)+ mRNA-seq data to minimize the impact of potential tsRNAs, additional experiments to assess whether these micropeptides are translation products of circRNA themselves remain necessary particularly when concerning with the mechanistic aspect of their synthesis.

In addition, some limitations might exist in our current study. In particular, a high-resolution view of cellular translational heterogeneity and cell-type-specific translational regulation was obscured by our bulk-seq data. Single-cell sequencing can be used to analyse the differences in gene expression between individual cells, but regrettably, technologies enable genome-wide investigation of *in vivo* gene translation with subcodon resolution at the single-cell level remain unavailable to date.

Nevertheless, our bulk-seq data measure the average behaviour of gene expression from heterogeneous tissue samples, this tends to reduce the sparsity of values within the expression matrix, making the data’s parameters more rich and less susceptible to dropouts. Our dataset can serve as a valuable resource for future studies of the translational machinery during retinogenesis, providing crucial and complementary information to single-cell omics data. In general, our present study provides an overall snapshot of gene translation and translational regulation during retinal development.

## DATA AVAILABILITY

All Ribo-Seq, RNA-Seq, and circRNA-seq sequencing data have been deposited in the SRA database at NCBI (https://www.ncbi.nlm.nih.gov/sra/) under accession number PRJNA589677. The mass spectrometry proteomics data have been deposited in the PRIDE database under accession number PXD023439.

## SUPPLEMENTARY MATERIALS

Supplementary materials can be found online.

## ACKNOWLEDGEMENT

We thank the support for the Center for Precision Medicine, Sun yat-sen University.

## AUTHOR CONTRIBUTIONS

Z.X. and H.W.W. supervised the project; J.Q.Y. performed RNA-seq and Ribo-seq library construction; C.Y.C. designed and performed experiments; M.Q.X provided help and suggestions for experimental design; K.N.C and H.H.L. analysed and interpreted the data; K.N.C., C.Y.C., H.W.W., and Z.X. wrote the manuscript; All authors approved the manuscript.

## REFERENCES

1. Bassett, E.A. and Wallace, V.A. (2012) Cell fate determination in the vertebrate retina. Trends in neurosciences, 35, 565–573.

2. Abdusselamoglu, M.D., Eroglu, E., Burkard, T.R. and Knoblich, J.A. (2019) The transcription factor odd-paired regulates temporal identity in transit-amplifying neural progenitors via an incoherent feed-forward loop. eLife, 8, e46566.

3. Lu, Y., Shiau, F., Yi, W., Lu, S., Wu, Q., Pearson, J.D., Kallman, A., Zhong, S., Hoang, T., Zuo, Z. et al. (2020) Single-Cell Analysis of Human Retina Identifies Evolutionarily Conserved and Species-Specific Mechanisms Controlling Development. Developmental cell, 53, 473–491.

4. Aldiri, I., Xu, B., Wang, L., Chen, X., Hiler, D., Griffiths, L., Valentine, M., Shirinifard, A., Thiagarajan, S., Sablauer, A. et al. (2017) The Dynamic Epigenetic Landscape of the Retina During Development, Reprogramming, and Tumorigenesis. Neuron, 94, 550–568.

5. Sze, Y.H., Zhao, Q., Cheung, J.K.W., Li, K.K., Tse, D.Y.Y., To, C.H. and Lam, T.C. (2021) High-pH reversed-phase fractionated neural retina proteome of normal growing C57BL/6 mouse. Scientific data, 8, 27.

6. Crick, F. (1970) Central dogma of molecular biology. Nature, 227, 561–563.

7. Kong, J. and Lasko, P. (2012) Translational control in cellular and developmental processes. Nature reviews. Genetics, 13, 383–394.

8. Teixeira, F.K. and Lehmann, R. (2019) Translational Control during Developmental Transitions. Cold Spring Harbor perspectives in biology, 11 (6), a032987.

9. Wang, H., Wang, Y., Yang, J., Zhao, Q., Tang, N., Chen, C., Li, H., Cheng, C., Xie, M., Yang, Y. et al. (2021) Tissue- and stage-specific landscape of the mouse translatome. Nucleic acids research, 49, 6165–6180.

10. Alvarez-Dominguez, J.R., Zhang, X. and Hu, W. (2017) Widespread and dynamic translational control of red blood cell development. Blood, 129, 619–629.

11. Atlasi, Y., Jafarnejad, S.M., Gkogkas, C.G., Vermeulen, M., Sonenberg, N. and Stunnenberg, H.G. (2020) The translational landscape of ground state pluripotency. Nature communications, 11, 1617.

12. Blair, J.D., Hockemeyer, D., Doudna, J.A., Bateup, H.S. and Floor, S.N. (2017) Widespread Translational Remodeling during Human Neuronal Differentiation. Cell reports, 21, 2005–2016.

13. de Klerk, E., Fokkema, I.F., Thiadens, K.A., Goeman, J.J., Palmblad, M., den Dunnen, J.T., von Lindern, M. and tHoen, P.A. (2015) Assessing the translational landscape of myogenic differentiation by ribosome profiling. Nucleic acids research, 43, 4408–4428.

14. Wang, Z.Y., Leushkin, E., Liechti, A., Ovchinnikova, S., Mößinger, K., Brüning, T., Rummel, C., Grützner, F., Cardoso-Moreira, M., Janich, P. et al. (2020) Transcriptome and translatome co-evolution in mammals. Nature, 588, 642–647.

15. Starr, C.R., Pitale, P.M. and Gorbatyuk, M. (2018) Translational attenuation and retinal degeneration in mice with an active integrated stress response. Cell death & disease, 9, 484.

16. Ji, Z., Song, R., Regev, A. and Struhl, K. (2015) Many lncRNAs, 5’UTRs, and pseudogenes are translated and some are likely to express functional proteins. eLife, 4, e08890.

17. Nelson, B.R., Makarewich, C.A., Anderson, D.M., Winders, B.R., Troupes, C.D., Wu, F., Reese, A.L., McAnally, J.R., Chen, X., Kavalali, E.T. et al. (2016) A peptide encoded by a transcript annotated as long noncoding RNA enhances SERCA activity in muscle. Science (New York, N.Y.), 351, 271–275.

18. Li, H., Xie, M., Wang, Y., Yang, L., Xie, Z. and Wang, H. (2021) riboCIRC: a comprehensive database of translatable circRNAs. Genome biology, 22, 79.

19. van Heesch, S., Witte, F., Schneider-Lunitz, V., Schulz, J.F., Adami, E., Faber, A.B., Kirchner, M., Maatz, H., Blachut, S., Sandmann, C.L. et al. (2019) The Translational Landscape of the Human Heart. Cell, 178, 242–260.

20. Martin, M. (2011) Cutadapt removes adapter sequences from high-throughput sequencing reads. EMBnet. journal, 17, 10–12.

21. Langmead, B., Trapnell, C., Pop, M. and Salzberg, S.L. (2009) Ultrafast and memory-efficient alignment of short DNA sequences to the human genome. Genome biology, 10, R25.

22. Liao, Y., Smyth, G.K. and Shi, W. (2019) The R package Rsubread is easier, faster, cheaper and better for alignment and quantification of RNA sequencing reads. Nucleic acids research, 47, e47.

23. Love, M.I., Huber, W. and Anders, S. (2014) Moderated estimation of fold change and dispersion for RNA-seq data with DESeq2. Genome biology, 15, 550.

24. Zhang, P., He, D., Xu, Y., Hou, J., Pan, B.F., Wang, Y., Liu, T., Davis, C.M., Ehli, E.A., Tan, L. et al. (2017) Genome-wide identification and differential analysis of translational initiation. Nature communications, 8, 1749.

25. Nueda, M.J., Tarazona, S. and Conesa, A. (2014) Next maSigPro: updating maSigPro bioconductor package for RNA-seq time series. Bioinformatics (Oxford, England), 30, 2598–2602.

26. Ingolia, N.T., Ghaemmaghami, S., Newman, J.R. and Weissman, J.S. (2009) Genome-wide analysis in vivo of translation with nucleotide resolution using ribosome profiling. Science, 324, 218–223.

27. Chothani, S., Adami, E., Ouyang, J.F., Viswanathan, S., Hubner, N., Cook, S.A., Schafer, S. and Rackham, O.J.L. (2019) deltaTE: Detection of Translationally Regulated Genes by Integrative Analysis of Ribo-seq and RNA-seq Data. Current protocols in molecular biology, 129, e108.

28. Gao, Y., Zhang, J. and Zhao, F. (2018) Circular RNA identification based on multiple seed matching. Briefings in bioinformatics, 19, 803–810.

29. Zhang, X.O., Dong, R., Zhang, Y., Zhang, J.L., Luo, Z., Zhang, J., Chen, L.L. and Yang, L. (2016) Diverse alternative back-splicing and alternative splicing landscape of circular RNAs. Genome research, 26, 1277–1287.

30. Zhang, J., Chen, S., Yang, J. and Zhao, F. (2020) Accurate quantification of circular RNAs identifies extensive circular isoform switching events. Nature communications, 11, 90.

31. Kim, D., Pertea, G., Trapnell, C., Pimentel, H., Kelley, R. and Salzberg, S.L. (2013) TopHat2: accurate alignment of transcriptomes in the presence of insertions, deletions and gene fusions. Genome biology, 14, R36.

32. Pamudurti, N.R., Bartok, O., Jens, M., Ashwal-Fluss, R., Stottmeister, C., Ruhe, L., Hanan, M., Wyler, E., Perez-Hernandez, D., Ramberger, E. et al. (2017) Translation of CircRNAs. Molecular cell, 66, 9–21.

33. Cox, J. and Mann, M. (2008) MaxQuant enables high peptide identification rates, individualized p.p.b.-range mass accuracies and proteome-wide protein quantification. Nature biotechnology, 26, 1367–1372.

34. Jones, P., Binns, D., Chang, H.Y., Fraser, M., Li, W., McAnulla, C., McWilliam, H., Maslen, J., Mitchell, A., Nuka, G. et al. (2014) InterProScan 5: genome-scale protein function classification. Bioinformatics, 30, 1236–1240.

35. Cepko, C. (2014) Intrinsically different retinal progenitor cells produce specific types of progeny. Nature reviews. Neuroscience, 15, 615–627.

36. Ren, Q., Yang, C.P., Liu, Z., Sugino, K., Mok, K., He, Y., Ito, M., Nern, A., Otsuna, H. and Lee, T. (2017) Stem Cell-Intrinsic, Seven-up-Triggered Temporal Factor Gradients Diversify Intermediate Neural Progenitors. Current biology, 27, 1303–1313.

37. Xiang, M. (2013) Intrinsic control of mammalian retinogenesis. Cellular and molecular life sciences, 70, 2519–2532.

38. Bicker, S., Khudayberdiev, S., Weiß, K., Zocher, K., Baumeister, S. and Schratt, G. (2013) The DEAH-box helicase DHX36 mediates dendritic localization of the neuronal precursor-microRNA-134. Genes & development, 27, 991–996.

39. Xu, X. and Pozzo-Miller, L. (2017) EEA1 restores homeostatic synaptic plasticity in hippocampal neurons from Rett syndrome mice. The Journal of physiology, 595, 5699–5712.

40. Bartesaghi, S., Betts-Henderson, J., Cain, K., Dinsdale, D., Zhou, X., Karlsson, A., Salomoni, P. and Nicotera, P. (2010) Loss of thymidine kinase 2 alters neuronal bioenergetics and leads to neurodegeneration. Human molecular genetics, 19, 1669–1677.

41. Ingolia, N.T., Lareau, L.F. and Weissman, J.S. (2011) Ribosome profiling of mouse embryonic stem cells reveals the complexity and dynamics of mammalian proteomes. Cell, 147, 789–802.

42. Wan, J., Masuda, T., Hackler, L., Jr., Torres, K.M., Merbs, S.L., Zack, D.J. and Qian, J. (2011) Dynamic usage of alternative splicing exons during mouse retina development. Nucleic acids research, 39, 7920–7930.

43. McNeely, K.C. and Dwyer, N.D. (2020) Cytokinesis and postabscission midbody remnants are regulated during mammalian brain development. Proceedings of the National Academy of Sciences of the United States of America, 117, 9584–9593.

44. Spriggs, K.A., Bushell, M. and Willis, A.E. (2010) Translational regulation of gene expression during conditions of cell stress. Molecular cell, 40, 228–237.

45. Johnstone, T.G., Bazzini, A.A. and Giraldez, A.J. (2016) Upstream ORFs are prevalent translational repressors in vertebrates. The EMBO journal, 35, 706–723.

46. Hinnebusch, A.G., Ivanov, I.P. and Sonenberg, N. (2016) Translational control by 5’-untranslated regions of eukaryotic mRNAs. Science, 352, 1413–1416.

47. Leppek, K., Das, R. and Barna, M. (2018) Functional 5’ UTR mRNA structures in eukaryotic translation regulation and how to find them. Nature reviews. Molecular cell biology, 19, 158–174.

48. Wu, Q., Wright, M., Gogol, M.M., Bradford, W.D., Zhang, N. and Bazzini, A.A. (2020) Translation of small downstream ORFs enhances translation of canonical main open reading frames. The EMBO journal, 39, e104763.

49. Minati, L., Firrito, C., Del Piano, A., Peretti, A., Sidoli, S., Peroni, D., Belli, R., Gandolfi, F., Romanel, A., Bernabo, P. et al. (2021) One-shot analysis of translated mammalian lncRNAs with AHARIBO. eLife, 10, e59303.

50. Szafron, L.M., Balcerak, A., Grzybowska, E.A., Pienkowska-Grela, B., Felisiak-Golabek, A., Podgorska, A., Kulesza, M., Nowak, N., Pomorski, P., Wysocki, J. et al. (2015) The Novel Gene CRNDE Encodes a Nuclear Peptide (CRNDEP) Which Is Overexpressed in Highly Proliferating Tissues. PloS one, 10, e0127475.

51. Schwinn, M.K., Machleidt, T., Zimmerman, K., Eggers, C.T., Dixon, A.S., Hurst, R., Hall, M.P., Encell, L.P., Binkowski, B.F. and Wood, K.V. (2018) CRISPR-Mediated Tagging of Endogenous Proteins with a Luminescent Peptide. ACS chemical biology, 13, 467–474.

52. Diallo, L.H., Tatin, F., David, F., Godet, A.C., Zamora, A., Prats, A.C., Garmy-Susini, B. and Lacazette, E. (2019) How are circRNAs translated by non-canonical initiation mechanisms? Biochimie, 164, 45–52.

53. Yang, Y., Fan, X., Mao, M., Song, X., Wu, P., Zhang, Y., Jin, Y., Yang, Y., Chen, L.L., Wang, Y. et al. (2017) Extensive translation of circular RNAs driven by N(6)-methyladenosine. Cell research, 27, 626–641.

54. Kristensen, L.S., Andersen, M.S., Stagsted, L.V.W., Ebbesen, K.K., Hansen, T.B. and Kjems, J. (2019) The biogenesis, biology and characterization of circular RNAs. Nature reviews. Genetics, 20, 675–691.

55. Wethmar, K., Barbosa-Silva, A., Andrade-Navarro, M.A. and Leutz, A. (2014) uORFdb--a comprehensive literature database on eukaryotic uORF biology. Nucleic acids research, 42, D60–67.

56. Li, J.J., Chew, G.L. and Biggin, M.D. (2019) Quantitative principles of cis-translational control by general mRNA sequence features in eukaryotes. Genome biology, 20, 162.

57. Chew, G.L., Pauli, A. and Schier, A.F. (2016) Conservation of uORF repressiveness and sequence features in mouse, human and zebrafish. Nature communications, 7, 11663.

58. Na, C.H., Barbhuiya, M.A., Kim, M.S., Verbruggen, S., Eacker, S.M., Pletnikova, O., Troncoso, J.C., Halushka, M.K., Menschaert, G., Overall, C.M. et al. (2018) Discovery of noncanonical translation initiation sites through mass spectrometric analysis of protein N termini. Genome research, 28, 25–36.

59. Renz, P.F., Valdivia-Francia, F. and Sendoel, A. (2020) Some like it translated: small ORFs in the 5’UTR. Experimental cell research, 396, 112229.

60. Starck, S.R., Tsai, J.C., Chen, K., Shodiya, M., Wang, L., Yahiro, K., Martins-Green, M., Shastri, N. and Walter, P. (2016) Translation from the 5’ untranslated region shapes the integrated stress response. Science, 351, aad3867.

61. Rapicavoli, N.A., Poth, E.M. and Blackshaw, S. (2010) The long noncoding RNA RNCR2 directs mouse retinal cell specification. BMC developmental biology, 10, 49.

62. Karali, M. and Banfi, S. (2019) Non-coding RNAs in retinal development and function. Human genetics, 138, 957–971.

63. Liang, W.C., Wong, C.W., Liang, P.P., Shi, M., Cao, Y., Rao, S.T., Tsui, S.K., Waye, M.M., Zhang, Q., Fu, W.M. et al. (2019) Translation of the circular RNA circβ-catenin promotes liver cancer cell growth through activation of the Wnt pathway. Genome biology, 20, 84.

64. Liu, Y., Li, Z., Zhang, M., Zhou, H., Wu, X., Zhong, J., Xiao, F., Huang, N., Yang, X., Zeng, R. et al. (2020) Rolling-translated EGFR Variants Sustain EGFR Signaling and Promote Glioblastoma Tumorigenicity. Neuro-oncology, 23(5), 743–756.

65. Ho-Xuan, H., Glažar, P., Latini, C., Heizler, K., Haase, J., Hett, R., Anders, M., Weichmann, F., Bruckmann, A., Van den Berg, D. et al. (2020) Comprehensive analysis of translation from overexpressed circular RNAs reveals pervasive translation from linear transcripts. Nucleic acids research, 48, 10368–10382.

